# Prioritizing working memory resources depends on prefrontal cortex

**DOI:** 10.1101/2024.05.11.593696

**Authors:** Grace E. Hallenbeck, Nathan Tardiff, Thomas C. Sprague, Clayton E. Curtis

## Abstract

How the prefrontal cortex contributes to working memory remains controversial, as theories differ in their emphasis on its role in storing memories versus controlling their content. To adjudicate between these competing ideas, we tested how perturbations to the human (both sexes) lateral prefrontal cortex impact the storage and control aspects of working memory during a task that requires human subjects to allocate resources to memory items based on their behavioral priority. Our computational model made a strong prediction that disruption of this control process would counterintuitively improve memory for low-priority items. Remarkably, transcranial magnetic stimulation of retinotopically-defined superior precentral sulcus, but not intraparietal sulcus, unbalanced the prioritization of resources, improving memory for low-priority items as predicted by the model. Therefore, these results provide direct causal support for models in which the prefrontal cortex controls the allocation of resources that support working memory, rather than simply storing the features of memoranda.

**SIGNIFICANCE STATEMENT:** Although higher-order cognition depends on working memory, the resources that support our memory are severely limited in capacity. To mitigate this limitation, we allocate memory resources according to the behavioral relevance of items. Nonetheless, the neural basis of these abilities remain unclear. Here, we tested the hypothesis that a region in lateral prefrontal cortex controls prioritization in working memory. Indeed, perturbing this region with transcranial magnetic stimulation disrupted the prioritization of working memory resources. Our results provide causal evidence for the hypothesis that prefrontal cortex primarily controls the allocation of memory resources, rather than storing the contents of working memory.

## INTRODUCTION

Working memory refers to our ability to both briefly store and perform operations on information no longer present. Our highest cognitive abilities depend on its function (Daneman and Carpenter, 1980; Engle et al., 1999), while its dysfunction cascades into a variety of cognitive symptoms characteristic of psychiatric disease (Silver et al., 2003). Decades of evidence (Funahashi et al., 1989; Miller et al., 1996; Courtney et al., 1998; Srimal and Curtis, 2008) evolved into mature theories that detail how feature-selective activity persists in populations of neurons in the dorsolateral prefrontal cortex (Goldman-Rakic, 1990; Compte et al., 2000), thus providing a neural mechanism for working memory storage. However, very little progress has been made in understanding the neural substrates and mechanisms underlying the processes that act upon and control information stored in working memory. While challenging to study, these control processes, the *working* in working memory, are what distinguish working memory from more passive short-term memory storage (Miller and Cohen, 2001; Curtis and D’Esposito, 2003). For instance, one can prioritize the resources allocated to memoranda based on their behavioral relevance to mitigate the hallmark capacity limitations of short-term memory (Luck and Vogel, 1997; Bays, 2014; Klyszejko et al., 2014; Sprague et al., 2016; Yoo et al., 2018).

We recently found that trialwise variations in the amplitude of persistent blood-oxygen-level-dependent (BOLD) activity in a visual field map in the superior branch of the precentral sulcus (sPCS) (Jerde et al., 2012; Mackey et al., 2017) in dorsolateral prefrontal cortex predicted the relative prioritization of two items decoded from visual cortex (Li et al., 2024). Here, we causally test the hypothesis that sPCS controls how resources are allocated to items stored in working memory based on their behavioral relevance. To do so, we measured the impact that transcranial magnetic stimulation (TMS) to sPCS had on working memory performance while participants maintained in working memory two items with different levels of relevance. The prioritized item was twice as likely to be tested after the memory delay. Theoretically, a controller should allocate a greater proportion of limited resources to the more likely target (Yoo et al., 2018). This predicts improved memory for high-priority targets at the cost of worse memory for low-priority ones, a pattern consistent with the behavior of human subjects (Bays and Husain, 2008; Klyszejko et al., 2014; Emrich et al., 2017).

Past work demonstrated that TMS and lesions to sPCS impair visual working memory for single items (Mackey et al., 2016; Mackey and Curtis, 2017), consonant with the idea that disruptions to a brain region subserving a task will generally degrade performance. Based on our computational model (Yoo et al., 2018), however, we predicted that perturbing sPCS during our prioritization task would disrupt the allocation process and counterintuitively improve memory performance for the low-priority target. To preview, this is indeed what we found. We additionally demonstrated that this effect was specific to sPCS and was not found when TMS was applied to the intraparietal sulcus (IPS2), another area associated with working memory (Todd and Marois, 2004; Xu and Chun, 2006; Li and Curtis, 2023).

## MATERIALS AND METHODS

### Participants

Seventeen neurologically healthy human participants (9 female, 8 male; mean age: 26.5, range: 22–33) performed in the no TMS version of the experiment. All participants had normal to corrected normal vision and were screened for TMS eligibility and excluded from participation if they had any brain-related medical issues or were currently taking certain drugs (e.g., antidepressants, amphetamines, chemotherapy, etc.). All participants gave written, informed consent and were compensated $10 per no TMS session and $50 per TMS session. After the initial analyses of the no TMS condition, three participants were excluded from the TMS analyses for failure to show a priority effect (i.e., greater saccade error on the low-priority items compared to the high-priority items; see Figure 1D). All subsequent analyses were performed with the remaining N = 14 (9 female, 5 male) participants. Sample size was estimated based on similar previous TMS studies (Mackey and Curtis, 2017).

**Figure 1.**
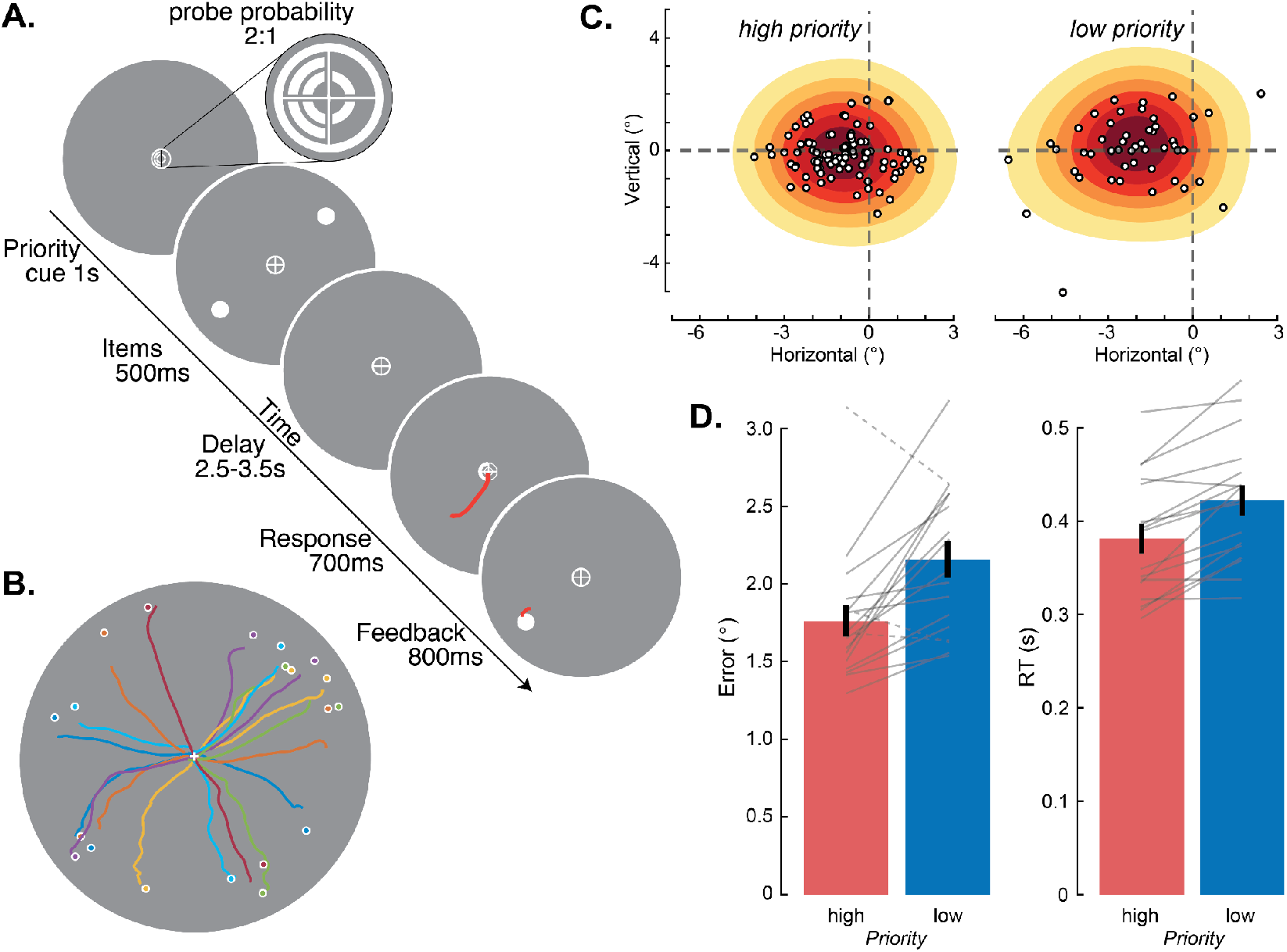
Item priority modulates memory error and saccade response times (no TMS). **A**, Memory-guided saccade task with two differentially-prioritized items. A precue indicated the priority of forthcoming items in each visual hemifield. Participants knew and were trained that high-priority items were probed twice as often as low items. After a retention interval, a response cue instructed which item should be the goal of the memory-guided saccade. Feedback was given, followed by an intertrial interval (2,000-3,000 ms; not shown). **B**, Saccade trajectories of single trials (colored lines) from an example participant, initiated from central fixation. Memory errors are defined as the Euclidean distance between endpoints of saccades and true item locations (circles). **C**, Distribution of memory errors for high-and low-priority items for an example participant. All items were rotated to a single polar angle from the origin (rightward). Note how memory errors for low-priority items were greater and the saccade endpoint distribution was less precise. Contours depict the empirical distribution over saccade endpoints, computed via kernel density estimation. **D**, At the group (N = 17) level, memory errors (left) were significantly lower (*t*(16) = −3.609, *p* = 0.003) and response times (RT; right) were significantly faster (*t*(16) = −5.415, *p* < 0.001) for high-priority compared to low-priority items. Error bars are standard errors of the mean (SEM). Lines denote data from individual participants. Dashed lines correspond to participants who were later excluded from the TMS study for failing to prioritize high-priority items (N = 3). Data are from the right visual hemifield, to match the data used in the analysis of TMS effects. Corresponding effects of priority were found in the left hemifield (Extended Data Figure 1-1).

### Experimental Procedures

Participants were seated 56 cm from the stimulus presentation monitor with their heads supported by a chin rest, which minimized movement during the task. Participants completed a two-item memory-guided saccade task (Figure 1A). The priority of the two items was established by manipulating which item was more likely to be probed for response after the delay, where the high-priority item was probed twice as often as the low-priority item. After participants attained fixation, a priority cue was displayed centrally, within the fixation crosshairs (1,000 ms). The priority cue indicated which half of the visual field (left or right) would contain the high-priority item, and which would contain the low-priority item. Subsequently, two working memory items (small white dots subtending 0.25°) appeared in the periphery (500 ms). The position of the items was previously determined such that one occupied the left hemifield and one occupied the right on every trial. After a delay period (2,500–3,500 ms, jittered), a response cue (half-circle, 700 ms) appeared at fixation, indicating which of the two items was the goal of a memory-guided saccade. Feedback was then provided by redisplaying the probed memory item (800 ms) and having participants make a corrective saccade to this location. An intertrial interval (ITI) then followed (2,000–3,000 ms). Participants were instructed to maintain fixation at the center of the screen during each trial, except when directed by the response cue to make a memory-guided saccade to the position of the cued item. They performed 36 trials/run, and completed 9 runs of the task on average (range: 4–17 runs) per TMS condition (no TMS, TMS to sPCS, TMS to IPS2).

### Transcranial Magnetic Stimulation (TMS)

We administered TMS using a 70 mm figure-eight air film coil (The Magstim Company, UK). The coil was positioned using the Brainsight frameless stereotaxic neuronavigation system (Brainsight, Rogue Research) and guided by a reconstructed T1 anatomical brain image. The coil was positioned tangentially on the scalp, with the coil parallel to the target ROI (i.e., with the coil handle perpendicularly bisecting the principal axis of the target region). TMS was applied in 3 pulses at 50Hz in the middle of the delay period of every trial, which in previous studies produced reliable effects across participants (Mackey and Curtis, 2017). Applying TMS to only one hemisphere confined the effect to the opposing visual field and consequently one memory item at a time, since one memory item occupied each hemifield. To limit the total number of TMS pulses in a day, the TMS condition was conducted over two sessions.

Given intersubject variability, including but not limited to cortical excitability and scalp-to-cortex distance (Kozel et al., 2000), we calibrated the TMS stimulator output per participant by measuring resting motor threshold (rMT) in a separate session, prior to the experimental TMS sessions. Motor threshold is stable over time, such that it can be used as a basis to target individualized protocols (Carpenter et al., 2012). To determine motor threshold, the coil was positioned 45° to the midsagittal plane on the precentral gyrus, and stimulation was delivered starting at 50% maximal stimulator output (MSO). This was modulated in 5% increments until visual twitches of the first dorsal interosseous muscle were evoked consistently, at which time MSO was steadily decreased until twitches were evoked either 3/6 or 5/10 times. rMT varied between 52% and 72% MSO across participants. During experimental sessions, we then applied TMS at 80% of each individual’s rMT.

We modeled the electrical field induced by TMS using SimNIBS (v. 3.2.6) (Thielscher et al., 2015). SimNIBS uses a segmentation of the anatomical scan along with the TMS parameters to compute the electrical field, taking into account the conductivities of different tissue types. We visualized the strength of the field for each participant in native anatomical space (Extended Data Figure 2-1 for sPCS and Extended Data Figure 4-1 for IPS2). To verify the spatial specificity of the stimulation, we extracted the average field strength from retinotopically-defined sPCS ROIs bilaterally (see Population receptive field mapping and definition of the sPCS below) and compared the field strength in the left sPCS TMS target to the field strength in right sPCS (Figure 2B).

**Figure 2.**
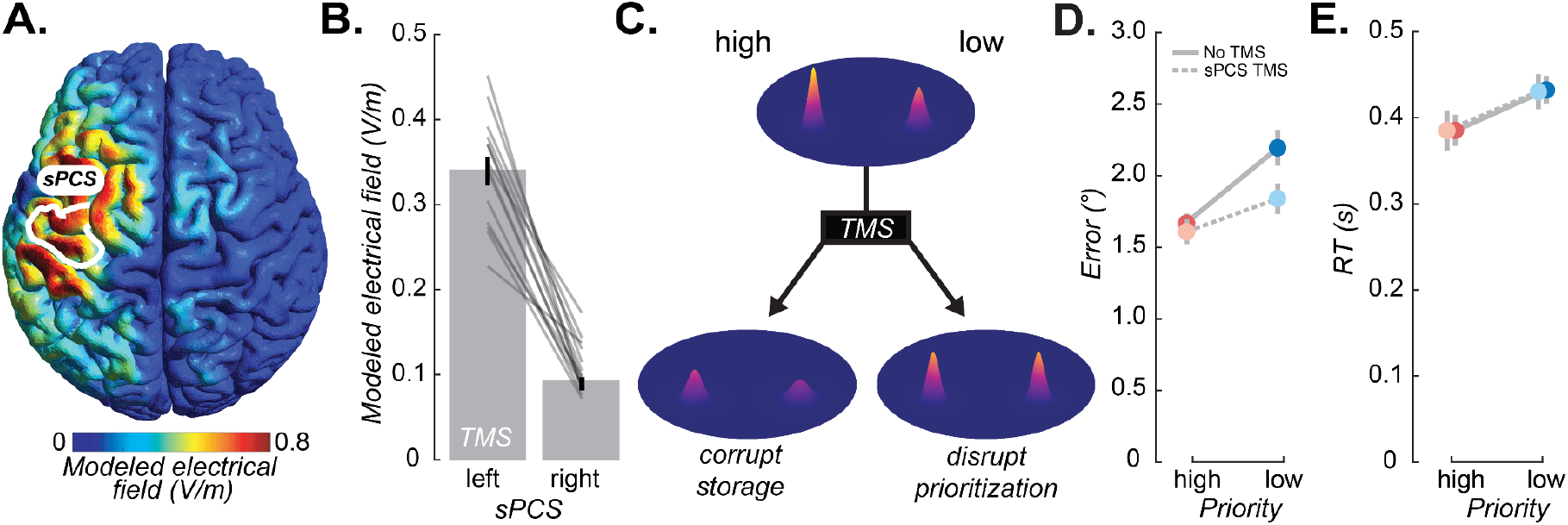
TMS to frontal cortex impacts working memory performance. **A**, The simulated electrical field induced by TMS targeting the retinotopically-defined left sPCS (white outline), for an example participant. See Extended Data Figure 2-1 for simulated electrical fields for all participants. **B**, The modeled electrical field strength in left (TMS) and right sPCS. Thin gray lines are individual participants (N = 14), bars are means across participants. Field strength is significantly greater in the left hemisphere, demonstrating the spatial specificity of the TMS (*t*(13) = 13.000, *p* < 0.001). **C**, Hypothesized effects of TMS on high- and low-priority items in working memory. We assume that memories are stored in noisy neural populations, where the gain of the neural activity corresponds to the precision of the memory (Ma et al. 2006). Without TMS (top), we hypothesized that the neural populations representing the high-priority item have a higher gain than those for the low-priority item. TMS could either corrupt the storage of the memoranda (bottom left) or it could disrupt the prioritization, such that the items are maintained with closer to equal gain (bottom right), leading to concomitant changes in memory error. Note that the relationship between changes in the precision of the representation and memory error are predicted to be nonlinear, and we present quantitative predictions in Figure 3. **D**, Mean (SEM) memory error plotted as a function of priority and TMS. Note how TMS lessened the difference between high- and low-priority items. **E**, Mean (SEM) saccade response times plotted as a function of priority and TMS. Data in **D**,**E** are for the contralesional hemifield. For ipsilesional results and individual participant data see Extended Data Figures 2-2, 2-3.

### Oculomotor procedures and analysis

Monocular tracking of gaze position was performed with the Eyelink 1000 (SR Research) recorded at 500 Hz. A 9-point calibration routine was performed at the start of each run. If, after multiple attempts, 9-point calibration failed, 5-point calibration was performed.

We preprocessed raw gaze data using custom software developed and regularly used by our lab (iEye, https://github.com/clayspacelab/iEye). This software implements an automated procedure to remove blinks, smooth the data (Gaussian kernel, 5 ms SD), and drift correct and calibrate each trial using epochs when it is known the eye is at fixation (delay) or the true item location (feedback). Memory-guided saccades were identified during the response period using a velocity threshold of 30 degrees/second. Trials were flagged for exclusion based on the following criteria: broken fixation during the delay; identified saccade < 2° in amplitude or > 150 ms in duration. Because TMS can often cause a facial flinch, including eyelid contraction, we removed 50 ms prior to and 150 ms following TMS pulses, ensuring any contraction-induced artifacts would not trigger the exclusion criteria. Overall, this resulted in usable data from 83% of trials on average, with a range of 54%–98% across participants. These procedures resulted in two behavioral outputs per trial: the endpoint of the memory-guided saccade and the initiation, or response time (RT) of the memory-guided saccade. We derived our primary behavioral measure—memory error—from the saccade endpoints by computing the Euclidean distance between the location of the saccade and the true location of the item.

### Statistical analysis

All statistical analyses were performed via permutation testing (10,000 samples). Repeated-measures analysis of variance was implemented using the permuco package (Frossard and

Renaud, 2021) in R (R Core Team, 2023). We performed *t*-tests in Matlab (Mathworks) using the PERMUTOOLS package (Crosse et al., 2024). Post-hoc tests were corrected for multiple comparisons using the Tmax method (Blair et al., 1994).

### Variable-precision model

To better isolate the mechanisms underlying the behavioral effects of TMS, we fit a variant of the variable-precision (VP) model to participants’ memory error data (van den Berg et al., 2012). The VP model is well-validated and has previously been shown to account for load effects (van den Berg et al., 2012) and priority effects (Yoo et al., 2018) in working memory. The model assumes that the precision of working memory, *J*, is variable across trials and items, where *J* is gamma-distributed with mean *J* and scale parameter τ. This formulation entails that the Gamma precision distribution has a shape parameter 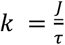 and and variance *J* τ. We modeled memory error, ε, as a Rayleigh distribution with parameter 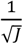, where ε is Euclidean distance, ε ≡ ‖**x**−**s**‖, **x** is the saccade endpoint, and **s** is the true item location. Therefore, the probability of error ε for a given precision distribution (e.g., in a given condition) is p(ε) = ∫ *Rayleigh*(*ε* | *J*)*Γ*(*J* | *J*, τ)*dJ*, with expected error E[ε] = ∫ *εp*(*ε*)*dε*. E[ε] is a nonlinear decreasing function of *J*, such that there is less change in error as *J* increases (Figure 3). Following Yoo et al. (2018) (Yoo et al., 2018), for the two-item priority task we assume that observers allocate a total amount of working memory resource, denoted *J*_*total*_, between the items in proportion to an allocation parameter, *εε*, such that the amount of resource allocated to the high priority item is *J*_*high*_ = *pJ*_*total*_ and the amount of resource allocated to the low-priority item is *J*_*low*_ = (1 − *p*)*J*_*total*_.

**Figure 3.**
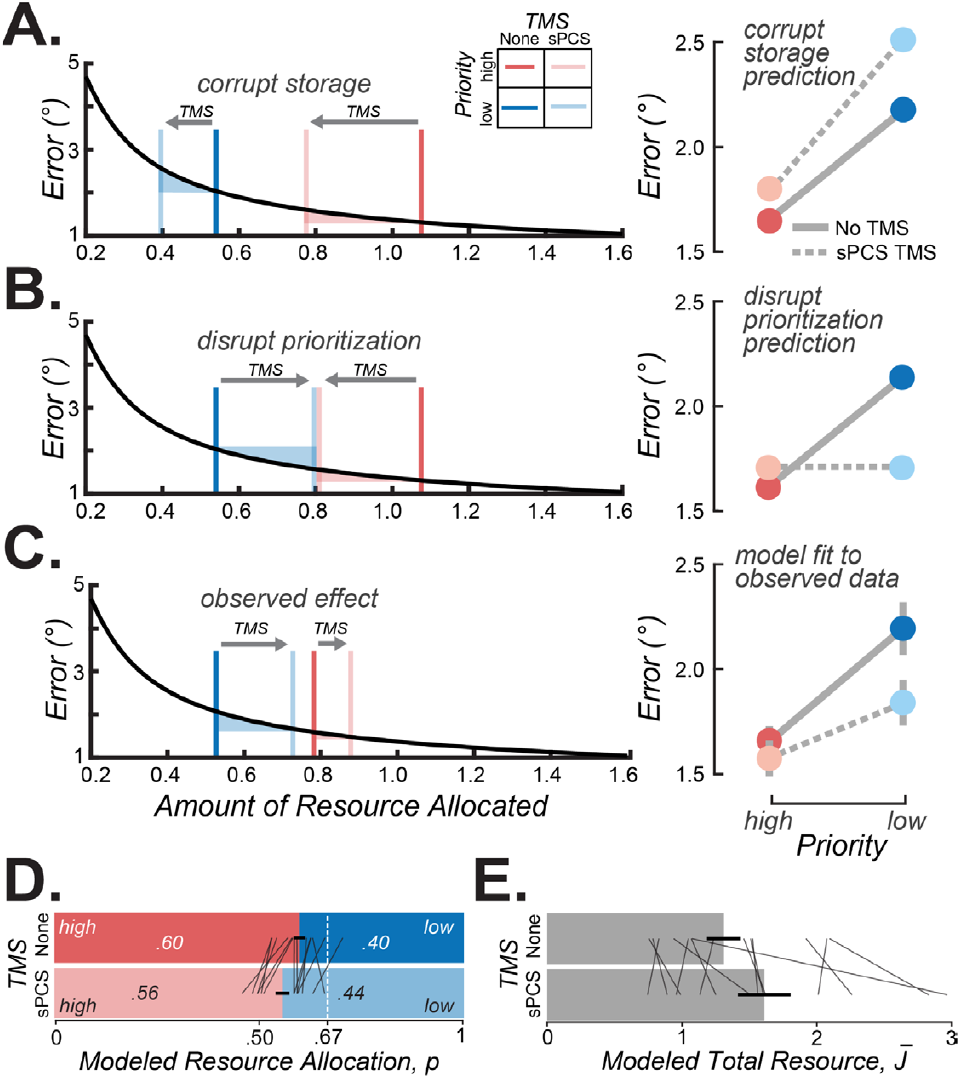
Predictions and fitted parameters from variable-precision model of working memory. **A**, In the model, the precision (inverse variance) with which each item is stored varies from trial-to-trial. The mean precision is controlled by parameter *J*. In the left plot, memory error decreases nonlinearly with increasing *J*, such that equal changes in *J* have a smaller effect on error at higher compared to lower precision, as depicted by the vertical component of the shaded regions between the bars. Assuming a pure storage corruption by TMS, the model predicts *J* parameters will shift leftward, resulting in greater memory error for both high- and low-priority items. The right panel plots the predicted mean memory errors if TMS had a pure effect on storage. **B**, Assuming a pure disruption of prioritization by TMS, each *J* for high- and low-priority would have an equal allocation (left). The right panel plots the predicted mean memory errors if TMS had a pure effect on prioritization. **C**, The average *J* derived from fitting the model to each participant’s data (left). The right panel plots the predicted mean memory errors from the model fits (gray solid and dashed lines) against the observed data (dots, means with SEM error bars, reproduced from Figure 2D). Note that the observed effects of TMS, which were well-captured by the model, are opposite in direction to the storage predictions for both low- and high-priority items. The effects match more closely the disrupted prioritization prediction. This was especially true for the low-priority item, which the model predicts will be most affected by TMS due to its lower precision. The range of the high-priority item in the no TMS condition may have been restricted by a floor effect (Extended Data Figure 3-1). **D**, The effect of TMS on the modeled *Resource Allocation* parameter (*p*) tested the possible disruptions of prioritization, as depicted in Figure 2C. The dashed white vertical line denotes the true probe probability for the high-priority item. **E**, The effect of TMS on the modeled *Total Resource* parameter (*J*) tested the possible corruption of working memory storage, as depicted in Figure 2C. Bars in **D** and **E** depict means (SEM error bars), and gray lines are individual participants.

**Figure 4.**
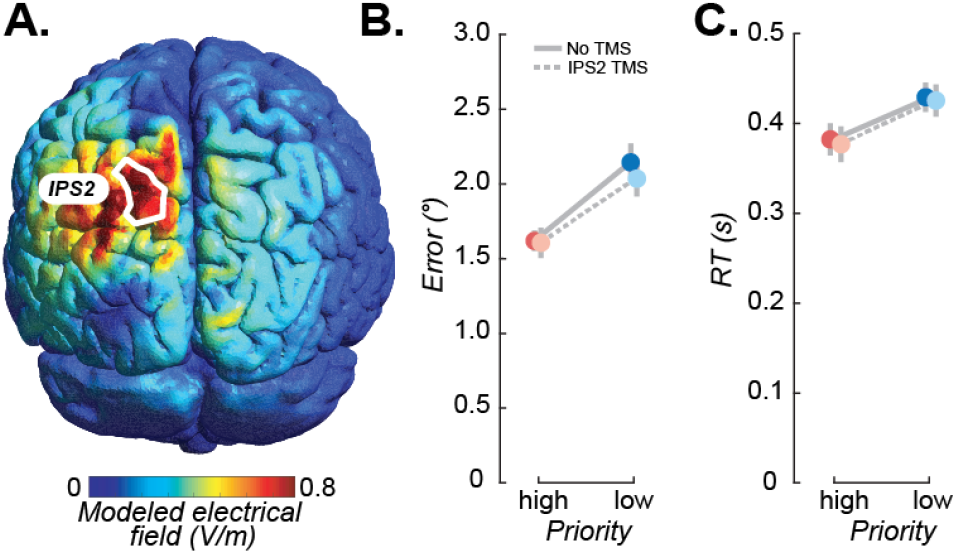
TMS to IPS2 had no effect on WM for items in contralesional hemisphere. **A**, The simulated electrical field induced by TMS targeting the retinotopically-defined left IPS2 (white outline), for an example participant. See Extended Data Figure 4-1 for simulated electrical fields for all participants. **B**, Memory error plotted as a function of priority and TMS. Points/thick lines: means across participants and error bars (SEM). Thin lines: individual participants. **C**, Saccade response times, plotted as in **B**. There were no differences between TMS conditions in error or RT (all *p*s > 0.05). For individual participant data see Extended Data Figures 4-2.

To assess the effects of TMS, we extended the model to allow *p* and *J*_*total*_ to vary between conditions, which were intended to model effects of TMS on prioritization and working memory storage, respectively. We fixed τ across conditions as we did not have any theoretically-motivated hypotheses about this parameter and did not have sufficient data to precisely estimate both the mean and scale of the precision distribution separately for each condition. As such, the model had five free parameters: *J*_*total,noTMS*_, *J*_*total,sPCS*_,*p*_*noTMS*_,*p*_*sPCS*_, τ.

We fit the parameters to each participant’s unaggregated, trial-level data using maximum-likelihood estimation in Matlab (fmincon). To avoid local minima, we fit the model 20 times per participant with different starting points for the optimization. Because there were approximately twice as much data for the high-priority condition as the low, we additionally randomly subsampled without replacement from the high-priority data on each iteration to match the amount of data for the low-priority condition, in order to avoid overfitting the model to the high-priority data. To generate behavioral predictions from the resulting model fits, we computed the prediction for the fit parameters from each model iteration and then averaged across the 20 iterations for display. We then averaged the parameter estimates across iterations prior to statistical analysis.

### Magnetic Resonance Imaging

Data were collected at New York University Center for Brain Imaging using a 3T Siemens Prisma MRI scanner (N = 14). Images were acquired using a Siemens 64-channel head/neck radiofrequency coil. Volumes were acquired using a T2*-sensitive echo planar imaging pulse sequence (repetition time (TR), 1200 ms; echo time (TE), 36 ms; flip angle, 66°; 56 slices; 2 mm x 2 mm x 2 mm voxels). High-resolution T1-weighted images (0.8 mm x 0.8 mm x 0.8 mm voxels) were collected at the end of the session, with the same slice prescriptions as for the functional data, and used for registration, segmentation, and display. Multiple distortion scans (TR 6,000 ms; TE 63.4 ms; flip angle, 90°; 56 slices; 2 mm x 2 mm x 2 mm voxels) were collected during each scanning session. The remaining three participants’ data were acquired using a 3T Siemens Allegra scanner using parameters described in (Mackey et al., 2017).

### Population receptive field mapping and definition of TMS targets

To define sPCS and IPS2, each participant underwent retinotopic mapping in the MRI scanner, following established procedures (Mackey et al., 2017). Participants maintained fixation at the screen center while covertly monitoring a bar aperture sweeping across the screen in discrete steps, oriented vertically or horizontally, depending on whether the sweep originated from the left or right or top or bottom of the screen, respectively. The bar was divided in thirds, with each segment containing a random dot kinematogram (RDK) used in a match-to-sample task. Participants reported which of the flanking RDKs moved in the same direction as the central RDK. Participants performed 8–12 runs of the task, with 12 bar sweeps per run. Task difficulty was staircased such that accuracy was maintained at 70–80%.

The resulting BOLD time series were fitted with a population receptive field (pRF) model with compressive spatial summation (Dumoulin and Wandell, 2008; Kay et al., 2013). We then identified left-hemisphere ROIs used as the TMS target on the basis of retinotopic and anatomical criteria. First, we visualized polar angle and eccentricity maps on the cortical surface, thresholded to include only voxels for which the pRF model explained > 10% of the variance. sPCS was then identified as the area at the junction of the superior prefrontal and precentral sulci containing a retinotopically organized representation of the contralateral visual field. Visual field maps in the intraparietal sulcus begin at the junction of the parietal-occipital sulcus and the intraparietal sulcus and proceed anteriorly, where the maps are demarcated by polar angle reversals. IPS2 is the third map along the sulcus and shares a foveal representation with IPS3.

### Data & Code Accessibility

All analysis and modeling code used in this work is available on a public GitHub repository: https://github.com/clayspace/{TBD}. The functionally-defined regions of interest, electrical field models, behavioral data, and modeling fits used here are available on the Open Science Framework: https://osf.io/{TBD}.

## RESULTS

We first replicated our previous behavioral results (Klyszejko et al., 2014; Yoo et al., 2018, 2022) showing that when given a precue that indicated the probability with which memory items would later be tested, memory errors were smaller and responses were faster for items that were more likely to be probed (Figure 1). Based on these behavioral measures, participants prioritized high over low-probability items in working memory.

We next asked if TMS to sPCS might disrupt the prioritization of memory resources. In human participants, we identified the sPCS using a modified population receptive field mapping technique with fMRI measurements (Mackey et al., 2017). We applied TMS to sPCS during the delay period and measured its effect on memory for items in the contralesional visual field (i.e., contralateral to the hemisphere where TMS was applied). We used a biophysical model of the electrical field induced by TMS, applied to each individual’s brain, which confirmed robust and accurate targeting of the left sPCS (Figure 2A,B; Extended Data Figure 2-1).

Our overall aim was to test two hypotheses regarding the mechanism by which sPCS supports working memory. First, if sPCS supports working memory storage, TMS should corrupt stored memories, resulting in a general increase in memory errors regardless of item priority (Figure 2C, left). Second, if sPCS controls the allocation of working memory resources, TMS should disrupt the process of prioritization, resulting in more similar memory errors across the two items (Figure 2C, right).

Focusing on the impact of TMS on memory errors for items in the contralesional visual field as a function of priority, a two-way repeated-measures ANOVA yielded main effects of priority (high, low) (*F*(1,13) = 22.366, *p* < 0.001), of TMS (none, sPCS) (*F*(1,13) = 8.545, *p* = 0.012), and a priority x TMS interaction (*F*(1,13) = 17.416, *p* = 0.001). TMS to sPCS significantly weakened the effect of priority compared to when no TMS was applied, driving the significant interaction (Figure 2D). Note the selective increase in accuracy following TMS for low-priority items (*t*(13) = 4.299, *p*_corrected_ *<* 0.001); TMS did not impact memory for the high-priority items (*p*_corrected_ > 0.05). TMS had no effect on the RT of memory-guided saccades (Figure 2E, all *p*s > 0.05). It also had no effect on working memory for items (all *p*s > 0.05) in the ipsilesional hemifield (Extended Data Figure 2-2; priority x TMS x hemifield interaction: *F*(1,13) = 5.809, *p* = 0.034), providing both an important control comparison and key evidence that TMS effects were spatially localized to the contralesional hemifield.

To provide additional insights into the extent to which TMS to sPCS affected storage versus allocation, we fit participants’ behavioral memory errors using our modified variable-precision model of working memory (Yoo et al., 2018). The model treats working memory as a continuous, noisy resource, such that across-trial variability in precision results from intrinsic variability in the amount of resource devoted to a given item (van den Berg et al., 2012). Importantly, our version of the model uses differences in memory precision as a function of item priority to estimate a parameter, *p*, that reflects the proportion of memory resources allocated to each item, as well as a parameter, *JJ*, that reflects the total amount of resource available for working memory (Yoo et al., 2018). These two model parameters map directly to the predicted effects of TMS depicted in Figure 2C. Critically, corrupted storage and disrupted prioritization make clear and divergent predictions about the effects of TMS on memory errors, as a function of the amount of resource allocated to the high- and low-priority items. However, the pattern of predicted memory errors from a pure corruption of storage (compare Figure 3A and 3C) were opposite in direction for both high- and low-priority items from what we observed. The pattern of predicted memory errors from a pure disruption of prioritization more closely matched the observed effects (compare Figure 3B and 3C).

Without TMS, the model confirmed that participants allocated significantly more resources to high-compared to low-priority items (*t*(13) = 7.430, *p* < 0.001, *H*_*0*_: *p* = 0.5). Moreover, replicating our previous results, the model demonstrated that without TMS participants overallocated resources to the low and underallocated resources to the high-priority items, relative to the objective probe probabilities used in the experiment (i.e., high:low 2:1) (*t*(13) = −5.173, *p* < 0.001, 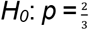), perhaps stemming from a strategy to minimize overall memory error (Yoo et al., 2018). In line with the disrupted prioritization hypothesis, TMS reduced the amount of working memory resource allocated to high-priority items, such that the resource was allocated more evenly between high- and low-priority items (*p* parameter; *t*(13) = −4.031, *p* = 0.002; Figure 3D). TMS also affected the total amount of working memory resource devoted to the two items (*J* parameter; t(13) = 2.029, *p* = 0.015; Figure 3E), resulting in a pattern of resource allocation and error that deviated somewhat from the pure predictions (Figure 3A,B). While the model predicts that changes in the high-priority item will be smaller than those for the low (i.e., note the very small vertical component of the pink shaded region between the lines denoting high-priority items in Figure 3A,B,C), these deviations may be partly explained by a floor effect for the high-priority item in the no TMS condition, limiting how small the errors could be. Indeed, when compared to a subset of participants for whom we measured working memory performance on a single-item version of the task, errors were statistically indistinguishable from errors for high-priority items (Extended Data Figure 3-1). Such a floor effect would then elicit an increase in the total resource parameter to offset the lack of change in the high-priority item. Despite the discrepancy, the overall pattern of observed effects better matched that predicted from a model in which TMS disrupted the allocation of working memory resource according to item priority.

While the above results spatially localize the effects of TMS and are qualitatively consistent with an effect on working memory prioritization, they do not establish whether this pattern of results is specific to sPCS, leaving open the possibility that TMS to other regions would elicit similar effects or alternatively would produce results more consistent with disrupting working memory storage (i.e., worse performance for both items). To control for this possibility, in the same subjects we additionally applied TMS to a region of parietal cortex (retinotopically-defined left intraparietal sulcus; IPS2, Figure 4A and Extended Data Figure 4-1 for simulated electrical fields of all participants), which is strongly implicated in working memory storage and control processes (Todd and Marois, 2004; Rahmati et al., 2018; Yoo et al., 2022), including suppressing information in working memory that is no longer relevant (Riddle et al., 2020). In contrast to sPCS, disrupting IPS2 had no effect on overall working memory performance or prioritization in the contralesional hemisphere compared to no TMS (all *p*s > 0.05; see Figure 4B/C and Extended Data Figure 4-2 for behavioral results for all participants). Therefore, these data indicate that the effects of TMS to sPCS were not due to a general effect of stimulation.

## DISCUSSION

We used a precise, retinotopically-guided TMS protocol to adjudicate between two competing hypotheses for the role of lateral prefrontal cortex in working memory, namely whether it is a substrate for working memory storage (Goldman-Rakic, 1990; Riley and Constantinidis, 2016) or instead that it exerts top-down control on storage regions in sensory cortex (Curtis and D’Esposito, 2003; D’Esposito and Postle, 2015; Serences, 2016; Tardiff and Curtis, 2024). During a two-item working memory task, TMS to sPCS selectively improved performance for the item with lower behavioral priority. Typically, disrupting a brain area impairs behavioral performance. However, TMS to sPCS did not cause a reduction in the quality of working memory. Instead, TMS disrupted a control process involved in the strategic allocation of the resources that support working memory. This disruption resulted in a more equal allocation of memory resources between the two items, despite their difference in behavioral relevance, and a striking improvement in performance for low-priority items. Our computational model of working memory provides a theoretical framework that explains how this selective improvement arises from the mechanisms by which the control of resources shapes the precision of working memory representations (Yoo et al., 2018). Furthermore, this disruption was specific to sPCS, as TMS to IPS2 did not affect working memory, and the effect of TMS to sPCS was hemifield-specific. These results provide clear and direct causal evidence that sPCS houses mechanisms critical for the control of working memory.

Evidence from previous research suggests that the sPCS, which is thought to contain the human homolog of the monkey frontal eye field (FEF) (Blanke et al., 2000; Mackey et al., 2017), may store spatial working memory representations. Robust and spatially-selective BOLD activity persists in human sPCS during working memory retention intervals (Courtney et al., 1998; Srimal and Curtis, 2008), and these patterns of activity can be used to decode item locations (Jerde et al., 2012; Hallenbeck et al., 2021; Li et al., 2021), albeit at a coarser level than from visual cortex. Moreover, surgical resections (Mackey et al., 2016) and TMS (Mackey and Curtis, 2017) to sPCS disrupt the accuracy of single-item memory-guided saccades. Parallel findings both from neurophysiology (Bruce and Goldberg, 1985; Funahashi et al., 1989) and inactivation (Sommer and Tehovnik, 1997; Dias and Segraves, 1999) studies of monkey FEF further establish the idea that FEF stores spatial working memory representations.

Our current results, however, invite a reconsideration of this interpretation. We hypothesize that instead of supporting a storage mechanism, the human sPCS prioritizes the allocation of resources that support working memory representations via top-down projections to cortical areas where these representations are stored. Abundant evidence instead points to visual cortex as the site of storage for visuospatial working memory (Serences, 2016; Curtis and Sprague, 2021; Dake and Curtis, 2024; Tardiff and Curtis, 2024). In a companion study, we used fMRI to decode high- and low-priority working memory representations while participants performed the same two-item working memory task used in the current study (Li et al., 2024). We found that trial-by-trial differences in the amplitude of persistent activity in sPCS predicted the relative quality with which the two prioritized items could be decoded from visual cortex. Specifically, when persistent activity was strong in sPCS during the delay, high-priority items were decoded from visual cortex with much better fidelity than low-priority items. Furthermore, there are known anatomical projections between monkey FEF and visual cortex (Stanton et al., 1995; Markov et al., 2014), and TMS to human sPCS modulates activity in visual cortex (Ruff et al., 2006). Microstimulation of FEF neurons improves behavioral performance during spatial attention (Moore and Fallah, 2001) and increases the gain of visually-evoked responses in V4 (Moore and Armstrong, 2003). FEF neurons projecting to V4 demonstrate strong, stimulus-selective delay-period activity during working memory, and the response properties of neurons in V4 and MT are modulated by the contents of working memory in a spatially-specific manner (Merrikhi et al., 2017). Together, these findings indicate that this region is well-positioned to exert top-down influences on visual cortex during both working memory and attention.

A caveat to this interpretation is the lack of effect of TMS on the high-priority item and the increase in the total resource parameter of the model. While this result could be due to a lack of independence of prioritization and storage processes at the neural level, two observations can explain these deviations. First, the model predicts that the impact of TMS on error will be very small for high-priority items because of the nonlinear relationship between total resource and error, consistent with our observation of little impact of TMS on high-priority items (Figure 3). Second, high-priority items in the no TMS condition may have been opposed by a floor effect, limiting how small the errors could be. Across all conditions of the current study, average error on the high priority item was approximately 1.6 dva, approximately the same as the memory error for single-item MGS data (i.e., with an effective priority of 1.0; Extended Data Figure 3-1). A combination of perceptual, oculomotor, and memory noise likely place an accuracy limit on working memory, preventing the high-priority items from further improvement in the no TMS condition. According to this account, the change in total resource is largely artifactual. Future work could mitigate this issue by using more difficult tasks (e.g. with more than two memoranda), which may be less susceptible to floor effects.

Furthermore, the idea that TMS to sPCS could lead to an increase in working memory storage capacity is contrary to a wealth of evidence that TMS to this region disrupts short-term memory and endogenous attention, including working memory (Mackey and Curtis, 2017), spatial priming (Campana et al., 2007; O’Shea et al., 2007) and trans-saccadic memory (Prime et al., 2010). With respect to attention, TMS to sPCS worsens performance on visual search tasks requiring top-down attention (Riddle et al., 2019) and diminishes validity effects induced by attentional cues, including reducing performance costs induced by invalid cues (Smith et al., 2005; Fernández et al., 2023). Considering that low-priority items are less likely to be tested, they resemble invalidly-cued items in studies of endogenous attention, suggesting sPCS may have a general role in maintaining a prioritized map of space (Jerde et al., 2012). The readout of such a map by visual cortex could bias processing in favor of neurons whose receptive fields match the prioritized portions of space, providing both a mechanism for the control of attention and working memory.

In summary, the present results provide direct causal support for the hypothesis that sPCS prioritizes information in memory. Such evidence is essential for moving beyond arguments over which of the many brain regions associated with working memory are essential to identifying the mechanistic role of each region and how they interact to support memory performance.

## Acknowledgments

We thank Mrugank Dake for assistance with TMS electrical field simulations.

## EXTENDED DATA

**Figure 1-1.**
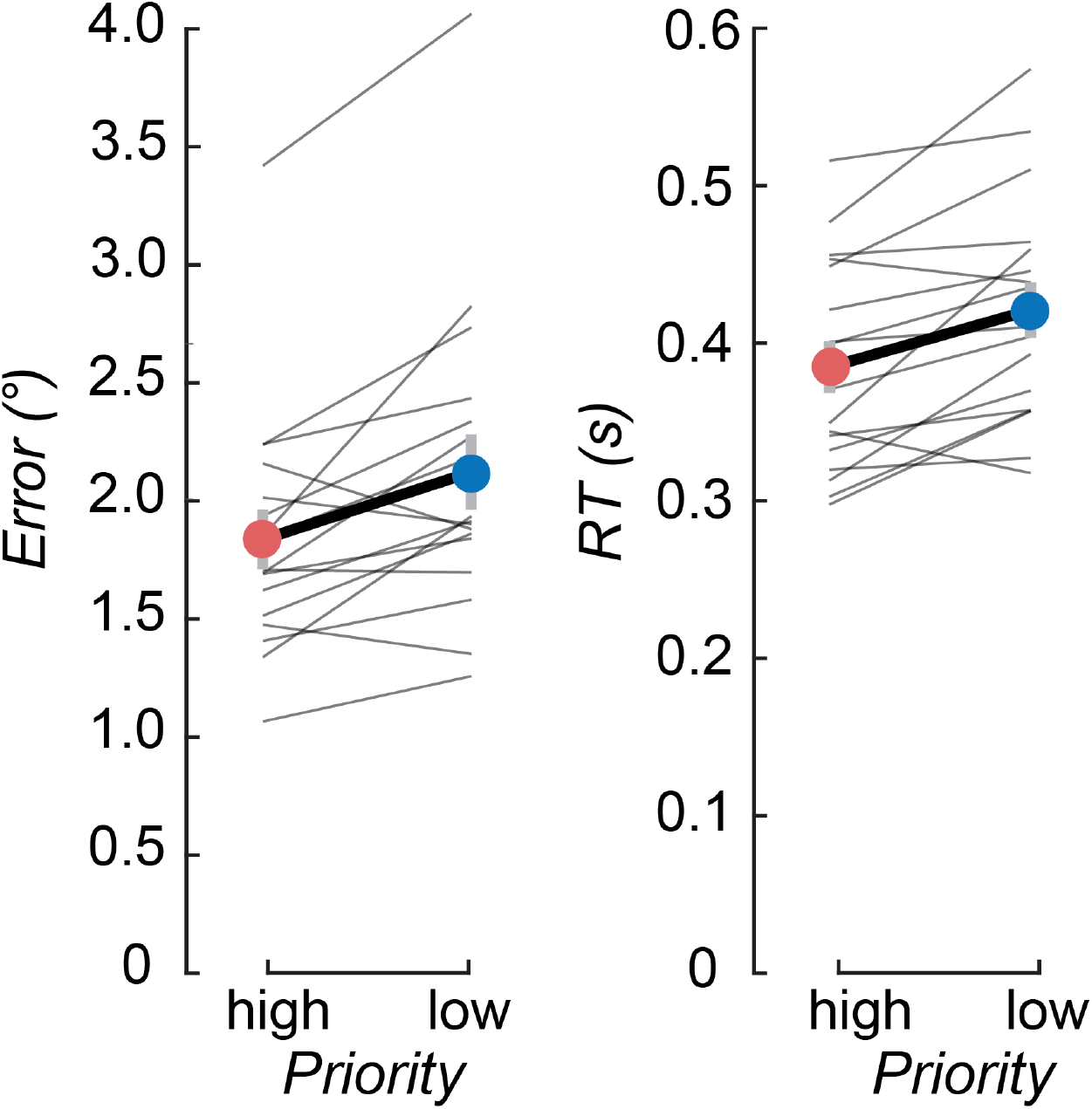
Memory error and saccade response times, left visual hemifield. Behavioral performance (N = 17): memory error (left; calculated as Euclidean distance from the item) and saccade RT (right) for items in the left hemifield. Errors were significantly lower (*t*(16) = −3.692, *p* = 0.002) and RTs were significantly faster (*t*(16) = −3.985, *p* < 0.001) for high-priority compared to low-priority items. Error bars are standard errors of the mean (SEM). Lines are data from individual participants.

**Figure 2-1.**
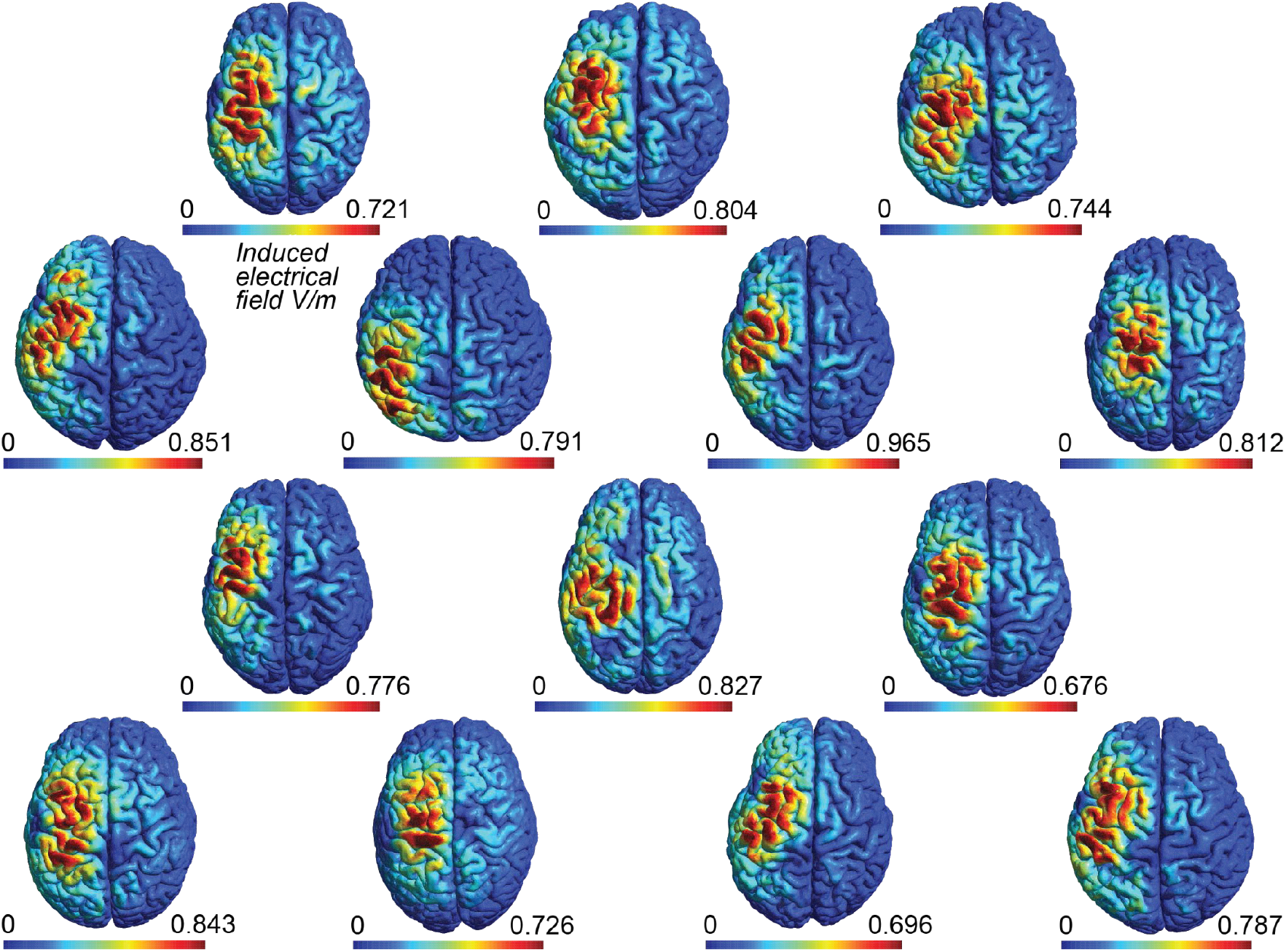
Simulated electrical fields induced by sPCS TMS. Simulated electrical field strength induced by TMS to left sPCS for all participants. See Online Methods for simulation details.

**Figure 2-2.**
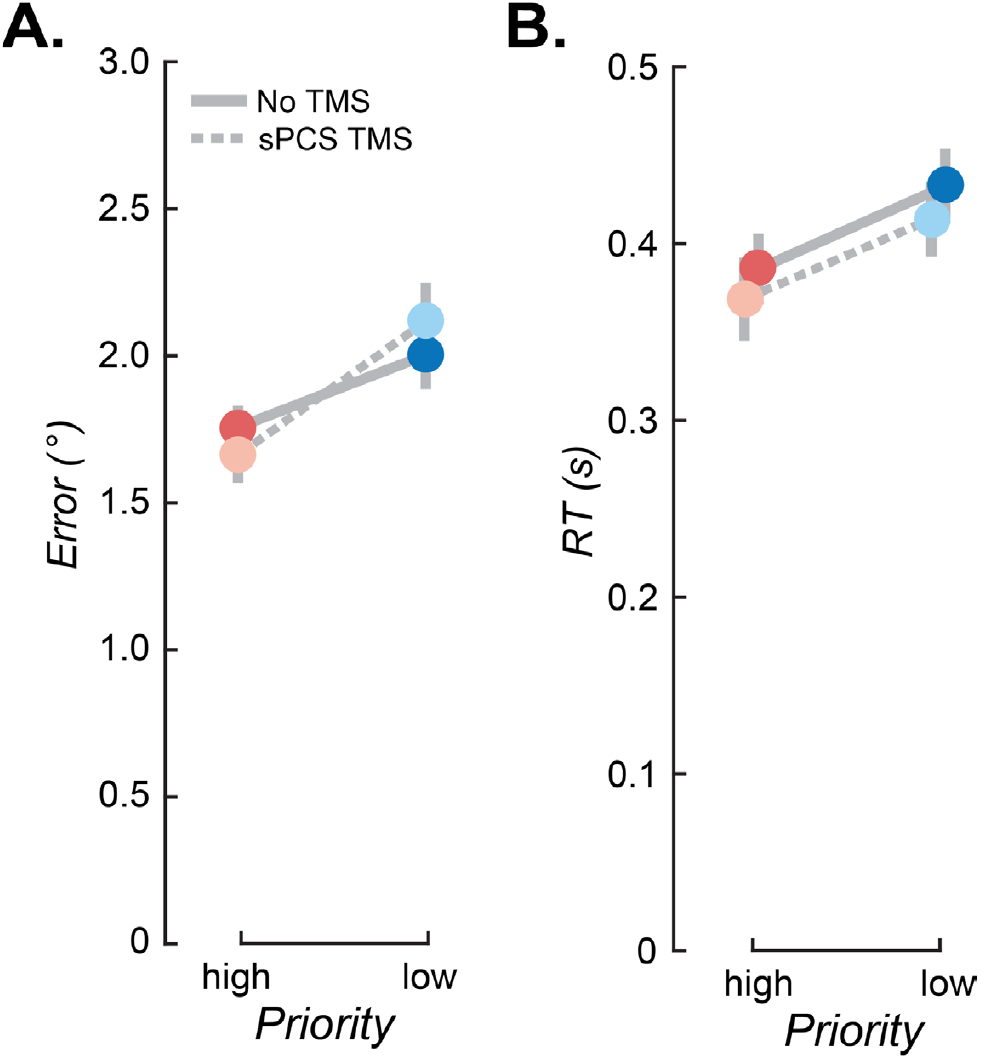
TMS to sPCS had no effect on WM for items in ipsilesional hemisphere. **A**, Memory error plotted as a function of priority and TMS. **B**, Saccade RTs plotted as in **A**. Error bars are SEM. There was no effect of TMS on errors or RT for items in the hemisphere ipsilateral to TMS (all *p*s > 0.05).

**Figure 2-3.**
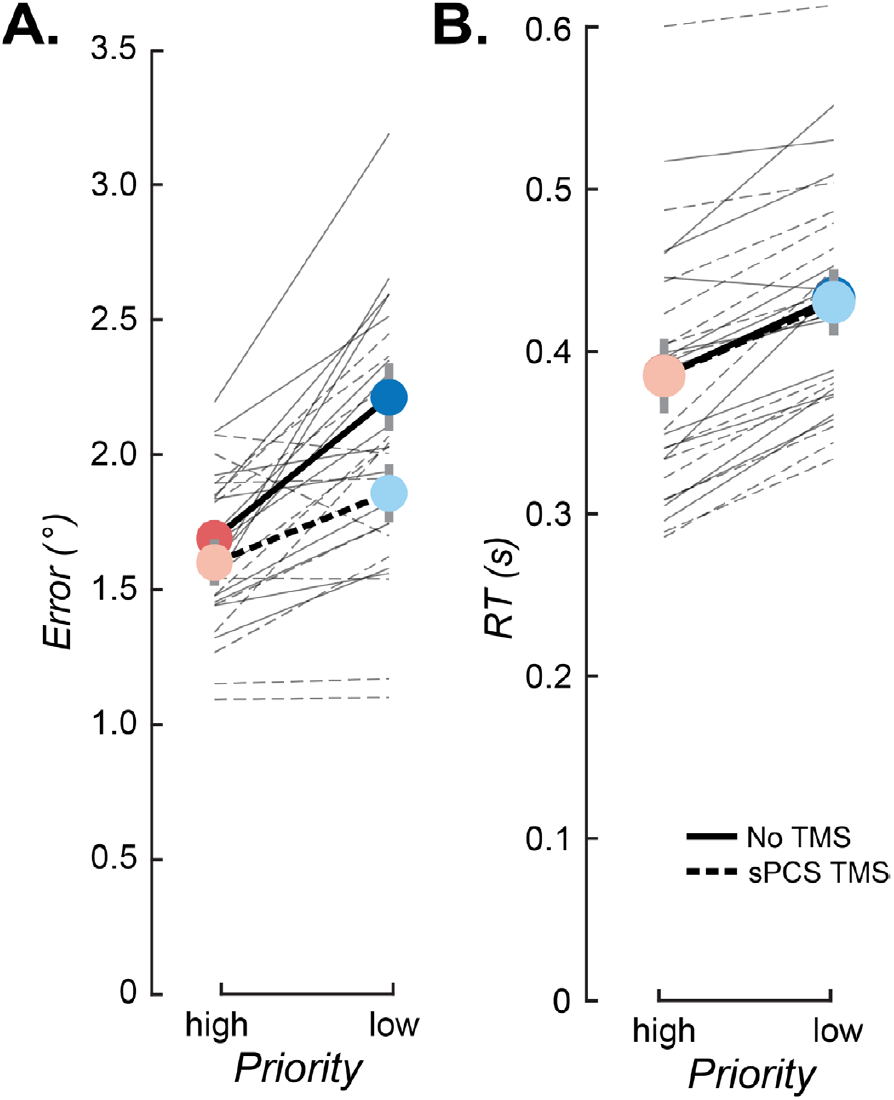
sPCS TMS effects on working memory performance for individual participants. **A**, Memory error plotted as a function of priority and TMS. Points/thick lines: means across participants and error bars (SEM), reproduced from Figure 3D. Thin lines: individual participants. **B**, Saccade response times, plotted as in **A**. Means and SEM reproduced from Figure 3E.

**Figure 3-1.**
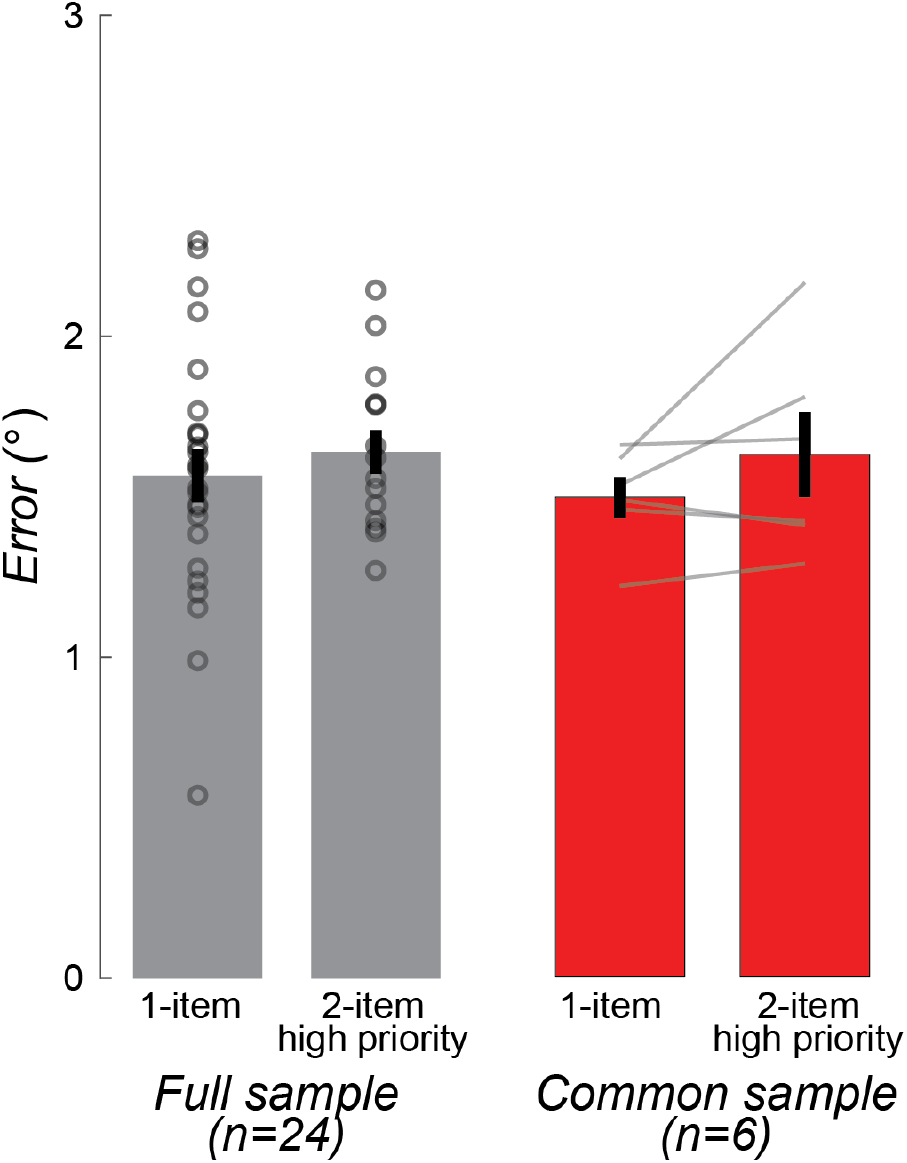
Errors for high-priority items do not differ from errors for a single item. Memory error as a function of memory load (one or two items). There was no difference in error between memory for a single item (N = 24) and memory for the high-priority item in the no TMS condition (N = 14) from the present study (gray; Mean difference (load 1 – 2) = –0.087°; *t*(36) = – 0.714, *p* > 0.05). Nor was there an appreciable difference within-participant for the subset of participants with data at both loads (N = 6, red; Mean difference = –0.146°; *t*(5) = –1.489, *p* > 0.05). Data points are individual participants. Error bars are SEM. Note that it is very rare for average memory error to fall below 1°, suggestive of a floor on WM precision. Load 1 data (N = 24) were compiled from three studies: Mackey & Curtis (2017)^1^ (N = 9) and two unpublished datasets (N = 17). All single-item studies had comparable memory delays (3–5 s) and general behavioral conditions to the present study. For participants with data in multiple single-item studies, errors were averaged over study before comparison (N = 2). All data are from the right hemifield to match the data in the TMS analysis.

**Figure 4-1.**
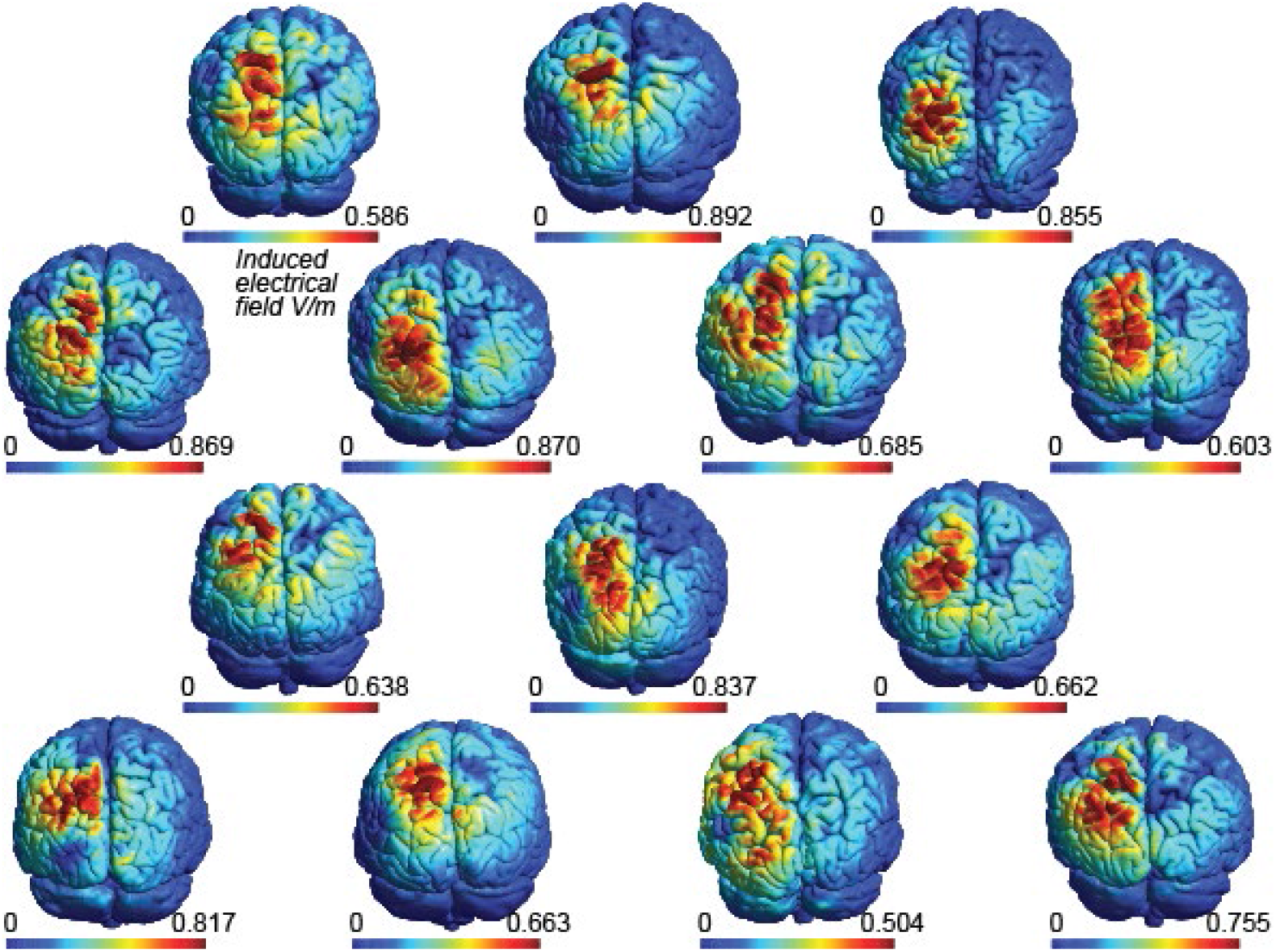
Simulated electrical fields induced by IPS TMS. Simulated electrical field strength induced by TMS to left IPS2 for all participants. See Online Methods for simulation details.

**Figure 4-2.**
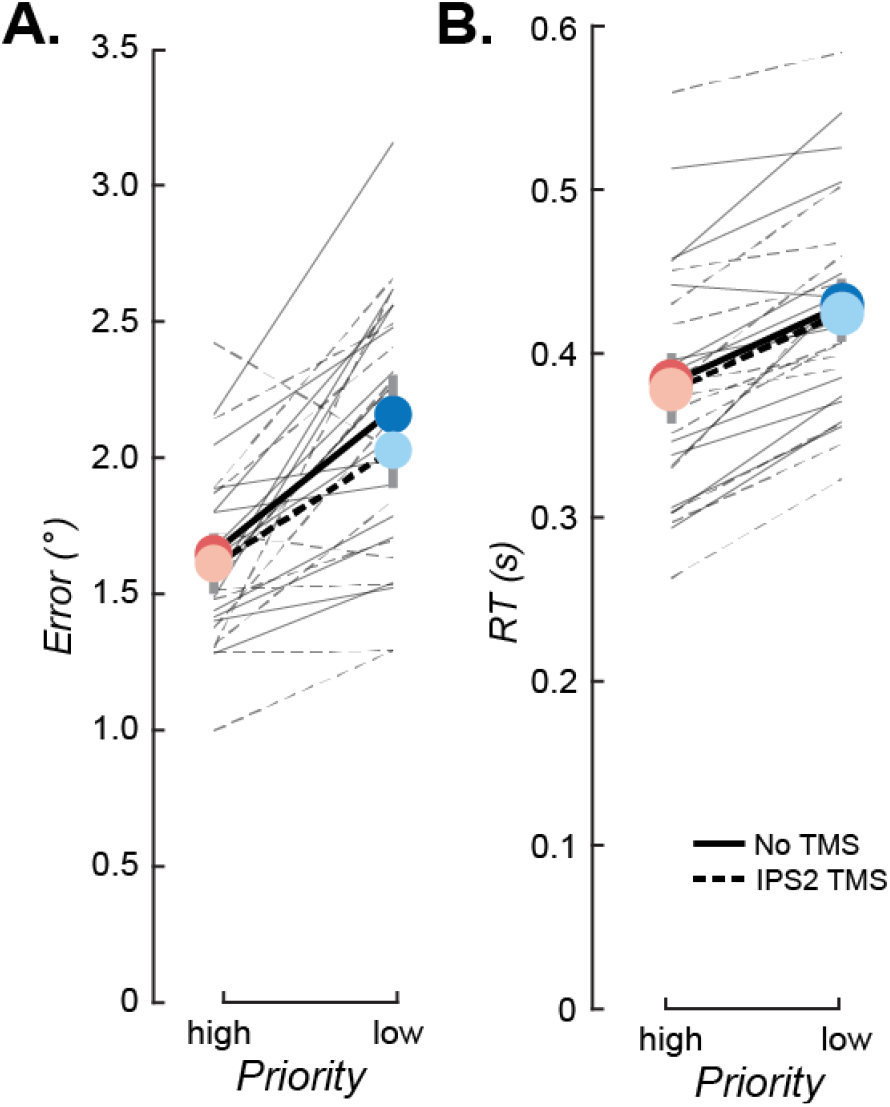
IPS TMS had no effect on WM for items in contralesional hemisphere. **A**, Memory error plotted as a function of priority and TMS. Points/thick lines: means across participants and error bars (SEM). Thin lines: individual participants. **B**, Saccade response times, plotted as in **A**. There were no differences between TMS conditions in error or RT (all *p*s > 0.05).

